# Exploiting node metadata to predict interactions in large networks using graph embedding and neural networks

**DOI:** 10.1101/2021.06.10.447991

**Authors:** Rogini Runghen, Daniel B Stouffer, Giulio V Dalla Riva

## Abstract

Collecting network interaction data is difficult. Non-exhaustive sampling and complex hidden processes often result in an incomplete data set. Thus, identifying potentially present but unobserved interactions is crucial both in understanding the structure of large scale data, and in predicting how previously unseen elements will interact. Recent studies in network analysis have shown that accounting for metadata (such as node attributes) can improve both our understanding of how nodes interact with one another, and the accuracy of link prediction. However, the dimension of the object we need to learn to predict interactions in a network grows quickly with the number of nodes. Therefore, it becomes computationally and conceptually challenging for large networks. Here, we present a new predictive procedure combining a graph embedding method with machine learning techniques to predict interactions on the base of nodes’ metadata. Graph embedding methods project the nodes of a network onto a—low dimensional—latent feature space. The position of the nodes in the latent feature space can then be used to predict interactions between nodes. Learning a mapping of the nodes’ metadata to their position in a latent feature space corresponds to a classic—and low dimensional—machine learning problem. In our current study we used the Random Dot Product Graph model to estimate the embedding of an observed network, and we tested different neural networks architectures to predict the position of nodes in the latent feature space. Flexible machine learning techniques to map the nodes onto their latent positions allow to account for multivariate and possibly complex nodes’ meta-data. To illustrate the utility of the proposed procedure, we apply it to a large dataset of tourist visits to destinations across New Zealand. We found that our procedure accurately predicts interactions for both existing nodes and nodes newly added to the network, while being computationally feasible even for very large networks. Overall, our study highlights that by exploiting the properties of a well understood statistical model for complex networks and combining it with standard machine learning techniques, we can simplify the link prediction problem when incorporating multivariate node metadata. Our procedure can be immediately applied to different types of networks, and to a wide variety of data from different systems. As such, both from a network science and data science perspective, our work offers a flexible and generalisable procedure for link prediction.

## Introduction

Real-world network datasets are often largely incomplete due to non-exhaustive sampling or the presence of complex hidden processes making data collection difficult [6,19]. As a result, accounting for incompleteness within data (e.g. “missing links”) is of key importance both to understand how different components in a system interact with one another and to accurately predict future trends in a system [8,13,24]. The action of predicting “missing links” or new links in a network is referred as link prediction in various fields [23,24]. The most commonly used methods to tackle link prediction include topological approaches, Block Model-based methods and graph-embedding methods. In topological methods, certain metrics describing the structure of a network (e.g. network properties such node degree and various centrality measures) are used to predict interactions [23]. Block Models-based approaches, such as the probabilistic generative family of Stochastic Block Models (and variants), aggregate nodes into groups based on their similarity of interactions [15,45]. Graph embedding methods on the other hand rely on projecting nodes onto an abstract latent feature space, so that the interaction probabilities depend on these latent features [2,5,43].

Multiple studies have shown that incorporating node metadata as covariates can both deepen our understanding of the network structure [30, 32, 33, 40], and improve link prediction accuracy in large scale networks [17, 26, 30]. However, incorporating node metadata presents various challenges. For instance, metadata diversity—i.e. whether the metadata variables are categorical or continuous—may require different modelling frameworks [3,26]. Due to the high number of nodes in large networks, they can be considered as high dimensional objects: indeed, when the network is represented as a matrix, each node is an additional coordinate. Therefore, accounting for metadata at the node level may also make the computational requirements overly demanding, as the complexity of the problem scales with the square of the number of nodes. To date, most attempts to incorporate node metadata for link prediction purposes have focused on node-aggregating methods such as Stochastic Block Models and its variants [17,26,40,41]. These methods make the prediction task more amenable by aggregating nodes into homogeneous groups. However, by doing so, they assume that all nodes within one group behave according to the same interaction probabilities, and thus are statistically indistinguishable [15, 45]. Unfortunately by disregarding the heterogeneity of interactions observed at the node level, such approaches oversimplify the network data. Here, we instead focus on using graph embedding methods which allows us to predict the interaction probabilities of each node directly, rather than aggregating the nodes in groups.

In our current study, we propose a new procedure that combines a graph embedding method with machine learning to predict interactions from nodes’ metadata. As the functional relationship between node metadata and the abstract latent feature spaces of a network is often unknown prior to data inspection, and can be very complicated, here we suggest using machine learning techniques to find an accurate mapping. In our procedure, we first use the graph embedding method to project nodes of the observed network on an abstract latent feature space at a lower dimensional space. By doing so, it allows us to learn a mapping from the nodes’ metadata to their abstract latent feature space (that we infer from the observed network) in an adequately low dimensional space. Because we move the problem from the original graph space to a lower dimensional feature space, our procedure simplifies the task of predicting interaction in large networks. Here, we specifically used neural networks as our machine learning technique to relate the observed nodes’ metadata onto the latent feature spaces of the observed network. The high flexibility of neural networks allowed us to account for the diversity of metadata. To illustrate the application of the proposed procedure in predicting interactions, we used a large dataset of tourist visits to destinations across New Zealand. Overall, our results showed that the proposed predictive procedure accurately predict interactions in large networks using both the knowledge from the observed network and the nodes’ metadata. Moreover, the proposed procedure also allowed us to predict interactions for new nodes better than at random.

## Materials and methods

In this article we focus on bipartite networks—i.e. networks that feature nodes of two types and with interactions (or links) that only occur between the different set of nodes. In the following sections, we describe: 1) the adopted network modelling procedure: first describing the Random Dot Product Graph model, then explaining how to infer, from an observed network, the position of its nodes in the latent space, and finally how to relate the nodes’ metadata to the nodes in their latent space using a machine technique; 2) an application to empirical network data; 3) the sensitivity and performance analyses we conducted to validate the proposed procedure.

### The Random Dot Product Graph model

Random Dot Product Graphs (RDPGs) are a class of Latent Position Models [14] developed to analyse social networks [31,47], and then extended to many other applications and types of networks [3,9,25,37,43]. To describe interactions in a network, such models assume that the probability of observing an interaction between two nodes is a function of the nodes’ features [31,47]. Here, we specifically use the RDPG implementation of Young and Scheinerman [47] to predict interactions in a bipartite context.

We define a bipartite network *G* as two distinct sets of nodes, *V* and *P* containing *M* and *N* nodes respectively, that is where *V* = *{v*_1_, …, *v*_*n*_*}* and *P* = *{p*_1_, …, *p*_*m*_*}* respectively; and a set of links, *E*, between the sets of nodes—i.e. (*v*_*i*_, *p*_*j*_) *∈ E*. We denote such a bipartite network as *G*(*V, P, E*). The bipartite network can be further represented as an adjacency matrix, where *G* is represented as the matrix *A ∈ {*0, 1*}*^*M ×N*^, where *A*_*ij*_ = 1 if (*v*_*i*_, *p*_*j*_) *∈ E* and *A*_*ij*_ = 0 otherwise.

In a bipartite RDPG model, each node *v*_*i*_ and *p*_*j*_ is assigned a vector of latent features *x*_*i*_ *∈* R^*d*^ and *y*_*j*_ *∈* R^*d*^. We call *d* the dimension of the network’s latent feature space, and the vectors *x*_*i*_ and *y*_*j*_ indicate the positions of the nodes *v*_*i*_ and *p*_*j*_, respectively, in the network’s latent feature space. The bipartite RDPG model further treats links as independent Bernoulli variables: two nodes interact with a probability equal to the dot product of their latent vectors, in formula:

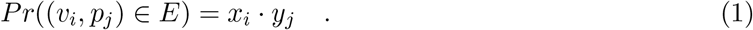

In matrix notation, we can represent the latent positions of all the nodes in *V* and *P* as the rows of a matrix **V** and the columns of a matrix **P**, respectively. As a result, the matrix of probabilities of interactions between the two node types in the network can be written as the matrix product **VP**.

For the matrix product to have meaning, the two matrices **V** and **P** need to have compatible dimension, which is satisfied if the latent feature spaces for *V* and *P* are equidimensional (that is, if the vectors *x*_*i*_ and *y*_*j*_ have the same number of coordinates). Moreover, for the matrix product to represent probabilities, the products must be in the [0, 1] range, which imposes additional geometric constraints in the latent feature spaces. Lastly, it is worth noting that any orthogonal transformation—e.g. a rotation—of **V** and **P** would result in an equivalent matrix of interaction probabilities: this will limit us to be able to infer the latent feature spaces up to an orthogonal transformation. Thus, we should refrain from reading any meaning in the absolute position of a node in the latent feature space.

### Inferring the position of nodes in the latent feature space

In theory, neither the nodes’ positions in the latent feature space, nor the dimension of the latent feature space are observable. Thus, we need to infer them from the observed network. To do so, we can exploit the adjacency spectral embedding—which is the truncated Singular Value Decomposition (SVD) of a network adjacency matrix—to obtain an unbiased estimate of the nodes’ positions in the network’s latent feature spaces [37].

The full rank SVD of an observed adjacency matrix *A* is given by three matrices *L*, Σ, and *R* such that *A* = *L ×* Σ *× R*^*T*^, with *L* and *R* real orthogonal matrices, and Σ a diagonal matrix whose entries are the singular values of *A* in decreasing order. As the sets of nodes *V* and *P* contain *M* and *N* nodes respectively, the matrices *L*, Σ, and *R* will have dimensions *M × S, S × S*, and *N × S*, respectively, where *S* = min (*M, N*). To compute the SVD of a matrix, we used the default svd function in R [35], which performed well for the large visitation data set described in later sections. (Note that fast algorithms that allow the decomposition of very large matrices such as in Liang et al. [22] and Zhou and Li [48] also exist if needed). We then used the profile-likelihood elbow criterion of Zhu and Ghodsi [49] to estimate an adequate dimension *d ≤ S* for the latent feature space.

Let *d* be the chosen dimension for the observed network latent feature space. We denote 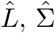, and 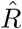 as the *d*-truncations of *L*, Σ, and *R*, respectively. We obtain them by retaining all the rows and the first *d* columns of *L* and *R*, and the first *d* rows and columns of Σ. We then compute the *d*-dimensional bipartite adjacency embedding of *A* as

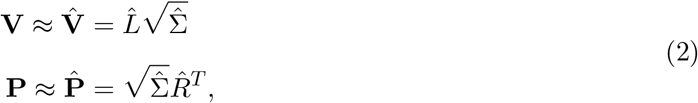

where 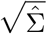 is a *d × d* diagonal matrix defined by the square root of the *d* greatest singular values of *A*.

### Predicting interactions using the nodes’ metadata via the latent feature space

Given the positions of all nodes in the latent feature spaces, the RDPG model completely determines the interaction probabilities between all nodes in the network. This implies that, if we were able to go from a node’s metadata to its position in the network’s latent feature space, we would be able to estimate its interaction probabilities with all other nodes in the network.

Let us consider the simplest scenario where we have a one-dimensional latent feature (in other words, the latent feature space is a vector with one coordinate, *d* = 1) and real valued metadata for the nodes in *V* and *P* . We can learn a mapping from the metadata to the latent space by fitting a linear regression model with the positions of nodes *v*_*i*_ in the latent feature space, *x*_*i*_ *∈* ℝ, as dependent variables and the metadata vectors *m*_*i*_ as independent predictor variables. Let *β*_0_ and *β* be the estimated intercept and vector of slope parameters for the linear model. The resulting model allows us to get the position of node *v*_*i*_ in the predicted latent feature space: *x*\*_*i*_ ∈ ℝ. We can then estimate the interaction probabilities of *v*_*i*_ via the dot product of *x*\*_*i*_ with the positions of the nodes in *P* ‘s latent feature space. Similarly, we can use the inferred linear model to predict the latent features of a *new* node added to the network—i.e. a node which was previously not observed *v*_*n*+1_—from the node’s metadata as followed: *x*_*n*+1_ = *β*_0_ + *m*_*n*+1_ *· β*. Then, to estimate the interaction probabilities of *v*_*n*+1_, we proceed with the dot product of *x*_*n*+1_ with the positions of the nodes in *P* ‘s latent feature space.

The latent feature space of large empirical networks is multivariate (even if not very large, 1 *< d « min*(*M, N*)). In general, nodes’ metadata are of different types—i.e. categorical and continuous. Finally, the relationship between metadata and latent features is often non-linear. Fortunately, a variety of statistical and machine learning approaches exist to solve the task of predicting a *d*-dimensional real valued vector from another vector (potentially larger and mixed valued). In particular, neural networks approaches can be used [12, 38]. In our application, we compare the performance of a classic linear regression, using ordinary least squares, and different neural network architectures.

To conclude, we have seen that we can use: a truncated SVD to estimate the latent feature spaces of nodes in a bipartite network, a variety of statistical and machine learning approaches to predict the latent features from the nodes’ metadata, and a simple dot product to predict interaction probabilities from latent features for nodes (Figure 1). In the next section, we apply our procedure to a large network of tourist-place visitation data deploying both linear models and two different neural network architectures.

**Figure 1:**
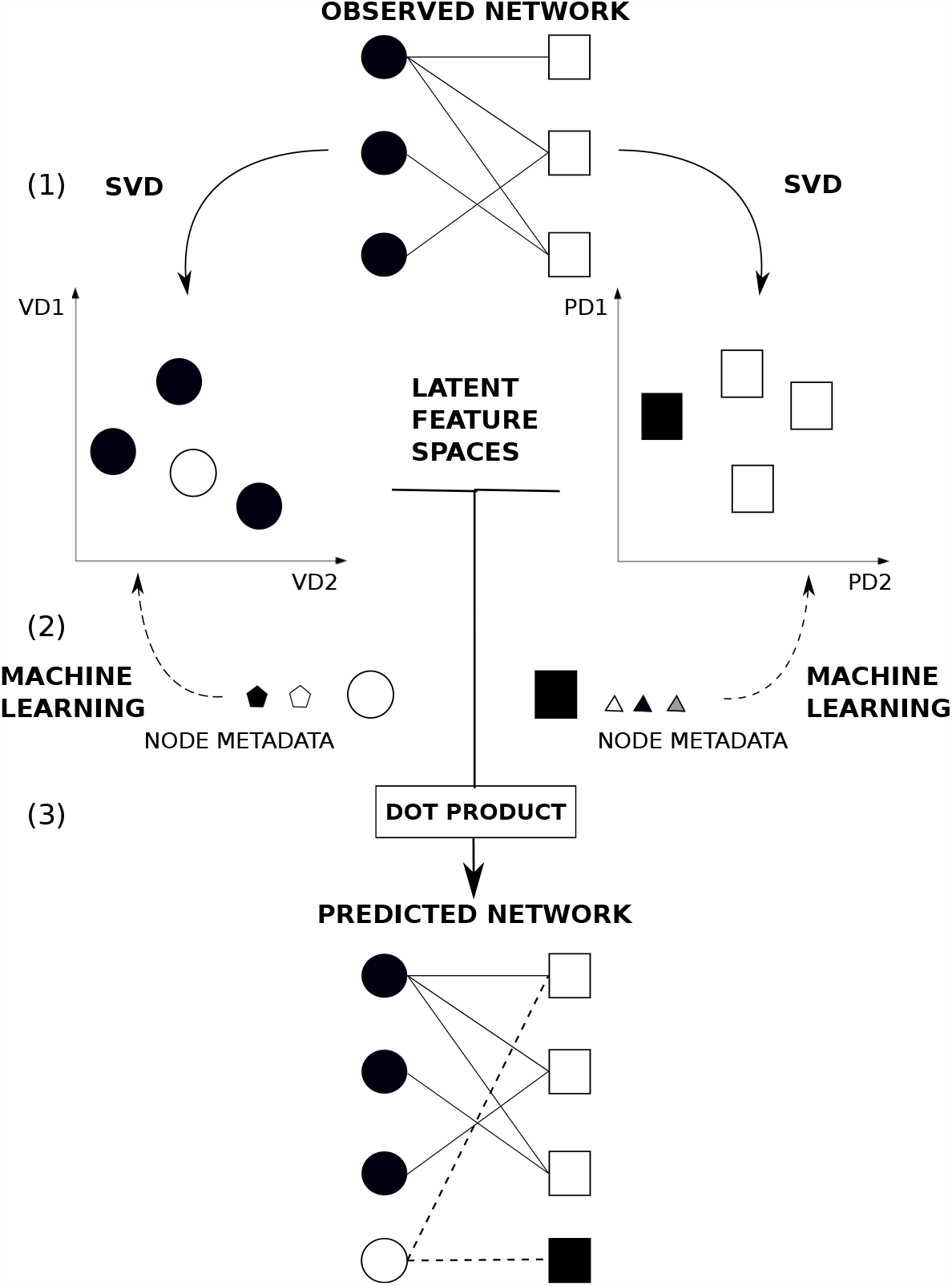
Using a Random Dot Product Graph Framework to predict interactions in a bipartite network. Note that here we use our case study—i.e. the travelling patterns of visitors to touristic destinations—to illustrate the framework. We first consider the travelling patterns of visitors as network data. In (1), the bipartite representation of travelling patterns of visitors: nodes are of two types—circles represent visitors and squares represent places, and links indicate a trip travelled by a given visitor to a given place. Here the solid lines represent the observed links. In (2), we first estimate the position of nodes within the observed bipartite network using a Singular Value Decomposition on the adjacency matrix representing the visitor– place interaction matrix. As a result, we obtain two latent feature spaces: a visitor latent feature space and place latent feature space. Note that here we show the embeddings of nodes of both visitors and places, respectively, in a latent feature space of dimension *d* = 2. We then relate the nodes’ metadata directly to their latent feature spaces. To do so, we use machine learning techniques to find the relationship between the two. To further predict *new* interactions in the network using observed metadata of new visitors and new places respectively—the models used to find the relationship of the metadata to the latent feature spaces are used. By doing so, we can project the new visitor and new place into the respective latent trait spaces LD1 and LD2. Finally, in (3), using the dot product, we are able to predict the probability of interaction between the new visitor and the new place added to the visitation network. Here, the dotted lines represent the new predicted links.

### Predicting tourist destinations from visitors and places metadata

To test the presented procedure, in this section we describe its application to visitation data representing the travel destinations of tourists across New Zealand. We specifically show how one can use visitor and place metadata, respectively, to estimate the interaction probabilities for visitors to travel to places within the country. We further show how one can potentially use the proposed procedure to predict *new* interactions—i.e. when new visitors and places are added to the visitation network—using only the respective nodes’ metadata and knowledge from the existing network.

#### Visitation data

To get an overview of the visitors’ travelling patterns across New Zealand, we extracted data from two national surveys conducted by the New Zealand Ministry of Business, Innovation and Employment (MBIE): The International Visitor Survey [28] and the Domestic Travel Survey [27]. The International Visitor Survey (IVS) targeted international visitors departing New Zealand at the 4 main international airports (Auckland, Wellington, Christchurch and Queenstown) whereas the Domestic Travel Survey (DTS) contacted domestic travellers via phone interviews about their recent trips. Both surveys record the list of places to which each visitor travelled during their trip within New Zealand. This accounted for a total of 189942 visitors travelling to 2616 places across the country. Note that these numbers refer to *only* visitor and places for which the complete set of metadata were available (Table 1 Supplementary Information).

**Table 1:** Summary of node metadata used

| Node type | Metadata | Data type | Classes |
| --- | --- | --- | --- |
| Visitor | Gender | Categorical | Male, Female |
| Visitor | Age | Categorical | Age group: < 20, 21–25, 26–34, 35–39, 40–44, 45–49, 50–51, 64–69, > 70 |
| Visitor | Activity type | Categorical | Hiking, Site-seeing, Water activities, Museums and other heritage sites, Visiting family, Work purposes |
| Visitor | Mode of transportation | Categorical | Car, Van, Boat, Tour bus, Bus, Helicopter, Aeroplane |
| Place | Place geolocation | Continuous | Latitude and Longitude of locations |
| Place | Place type | Categorical | Heritage site, Crown Protected area, Town, Village, Recreational site |
| Place | Regional Council | Categorical | Northland, Auckland, Waikato, Bay of Plenty, Gisborne, Hawke’s Bay, Taranaki, Manawatu-Wanganui, Wellington, Tasman/Nelson, Marlborough, West Coast, Canterbury, Otago, Southland, and Areas Outside Regional Council |

**Table 2:** Summary of number of visitors and places used for the different steps of the predictive procedure

| Dataset | Analysis | No. of visitors | No. of places |
| --- | --- | --- | --- |
| Full training set | SVD | 101656 | 430 |
| Model training set | Neural Network | 71159 | 301 |
| Model test set | Neural Network | 30497 | 129 |
| Validation set | Neural Network, Dot Product | 88286 (43662 *new*) | 360 (120 *new*) |

#### Node attributes: visitor and place metadata

Visitor metadata includes age, gender, activity type, and the mode of transportation used during their trip (Table 1). These characteristics are present for both the Domestic Travel Survey and the International Visitor Survey. As the survey data only had the name of places visited by travellers, we had to define the attributes of the different places within the visitation data. In general, places across New Zealand can be categorised based on the ownership of the land or the type of activities performed on those lands. Therefore, to identify the land type of each place within the visitation data, we used the geospatial maps provided by Land Information New Zealand [20] to map the places extracted from the different national surveys. For example by doing the latter, this allowed us to distinguish whether a particular place was categorised as a recreational site or national heritage site (Table 1).

#### Predicting visitor-place interactions in visitation network using node metadata

Here we are particularly interested to test our predictive procedure in two different contexts: predicting interaction probabilities of nodes present in the observed network, which we refer as “observed” nodes, and predicting interaction probabilities of nodes that we artificially removed from the original data set, which we refer as “new” nodes. Artificially removing nodes from the observed network allows us to further test the ability of the proposed predictive procedure to predict interaction probabilities of out-of-sample nodes—i.e. when either new visitors or places are added to the network—using their metadata. As such, we split the visitation data into a training and a validation set. The training set contains the *observed* nodes and validation set contains the *new* nodes. As the validation data serves as the *new* data, we made sure that the validation set contained both visitors and places identities not present in the training data set. Note that we build the observed visitation network from the interaction data contained in the training set and computed its adjacency matrix. In the rest of this section, we explain our predictive procedure in detail. The procedure involves three key steps: (1) we perform a Singular Value Decomposition of the adjacency matrix of the observed network to compute the position of the observed nodes (from the training data) in their latent feature space; (2) we use the training data to first fit regression models that predict the nodes’ positions in the latent feature space as a function of the nodes’ metadata; then, we use the fitted models to predict the position of the nodes from the validation data in their latent feature space; (3) we use these predicted positions to estimate the interaction probabilities of nodes in the validation set.

1. We computed the Singular Value Decomposition (SVD) of the training network’s adjacency matrix *A*_*T*_ . Then, truncating the SVD, we computed the positions of the observed visitor and place nodes in their respective latent feature spaces, 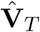 and 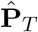 respectively:

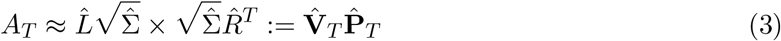
2. We fit three different types of multivariate regression models on the training data set. Let **v**_*T*_ and **p**_*T*_ be the metadata for visitor and places nodes in the training set, then the regression modelling task is to find a pair of function 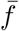 and 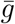 such that 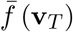 and 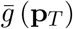best approximate 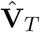 and 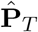 respectively, where 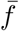 and 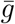 are part of some family of functions 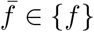 and 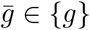. For sake of clarity, 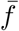 and 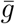 are function from the space of metadata (visitor nodes’ and place nodes’ metadata respectively) to the space of latent features (for visitors and places respectively). Namely, we fit: a) a linear regression—where we used a linear function to relate directly the metadata to the latent feature space (specified as in Table S3); b) a multilayer perceptron (MLP)— i.e. a neural network with one dense hidden layer of 200 nodes using a rectified linear unit (ReLU) as our activation function; and c) a neural network with two dense hidden layers (NN) of 250 nodes each and the ReLU activation function. Due to the variety of data types—i.e. varying from categorical to continuous variables (Table S2)—and the high flexibility of neural networks in solving regression and classification problems [16], we compared two learning rates: a constant learning rate of 0.01 and a time-based decay—where the initial learning rate (0.01) decreased by 0.0001 after each epoch. We used the Mean Absolute Error to measure the distance between the predicted and estimated latent features and assess the accuracy of the model training (refer to Table S4 to see the results obtained using other metrics). To monitor the training of the different models and ensure that they were not overfitting, we split the training data set into two sets: a training set (70 % of the training set) and a test set (30 % of the training set). We trained all the regression models on the training set and evaluated their accuracy on the test set. We used Google’s deep learning software TensorFlow [1] and Keras [7] implemented in Python 2.7 [44] to fit all the aforementioned models using the adaptive moment estimation (Adam) optimiser [18] with 30 epochs and a batch size of 20. We then used the fitted multivariate regression models to predict the positions of the nodes from the validation data set in the latent feature space, 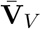 and 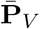 respectively. The predicted values 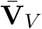 and 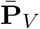 are functions of the nodes’ metadata:

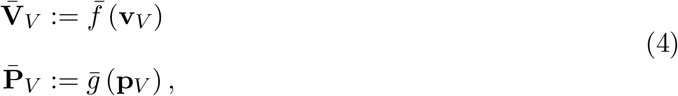

where **v**_*V*_ and **p**_*V*_ are the metadata for visitor and places nodes in the validation set, and 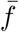 and 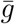 are the function obtained from the training data.
3. Using the nodes’ positions 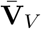 and 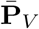 obtained by the models in (2), we estimated the interaction probabilities for all nodes present in the validation data and the nodes in the training data set by multiplying the matrices containing the nodes’ position in their respective latent feature spaces:

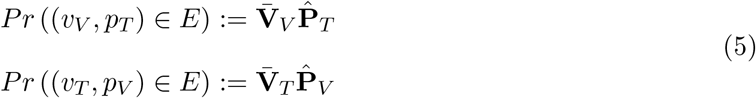

where, with some abuse of notation, *Pr* ((*v*_*V*_, *p*_*T*_) *∈ E*) and *Pr* ((*v*_*T*_, *p*_*V*_) *∈ E*) are the matrices of interaction probabilities between visitor nodes in the validation set and places nodes in the training set, and between visitor nodes in the training set and places nodes in the validation set.

Specific pairwise interaction probabilities can be estimated multiplying the vector of the predicted latent feature position of new nodes (inferred from the regression methods) to the latent features position vectors (estimated from the SVD) for all observed nodes present in the observed network, that is, using the dot product. For example, considering a new visitor node *n* with metadata **v**_*n*_, a predictive function 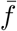, and a known place node *p*, whose position in the latent feature space (as obtained by SVD in step 1) is *x*_*p*_, the interaction probability between *n* and *p* is:

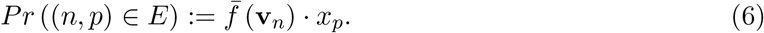

#### Sensitivity and performance analysis of predictive procedure

We calculated the probability of interaction between the nodes in the test set as:

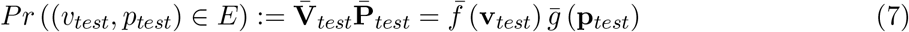

where *v*_*test*_ and *p*_*test*_ are the nodes in the test set, 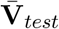 and 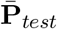 are the predicted latent features positions, **v**_*test*_ and **p**_*test*_ are the nodes metadata, and 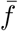 and 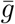 are the trained predictive functions. To assess the performance of the overall predictive procedure, we calculated the sensitivity—i.e. the ratio of correctly predicted links to observed links, and the accuracy—i.e. the ratio of correctly predicted observed links (True Positive) and correctly absent links (True Negative)—in our test data [42].

Furthermore, to evaluate and assess the performance of the different combinations of RDPG-regression models in correctly predicting the observed interactions, we used the Area Under Curve-Receiver Operator Curve (AUC-ROC) to evaluate the performance of the different combinations. To do so, we calculated the rate of True Positives—i.e. predicting an interaction when it is actually present—and False Positives—i.e. predicting an interaction when it is actually absent—at different thresholds varying from 0 to 1. AUC-ROC is used as a measure to assess the ability of different models to distinguish between a True Positive and a False Positive. For instance when 0.5 *< AUC ≤* 1, this indicates that the predictive model is performing well—i.e. the model effectively distinguishes a True Positive from a False Positive—whereas 0 *≤ AUC <* 0.5 indicates that the predictive model is not effectively distinguishing between True Positives and False Positives.

## Results

For the training visitation dataset (number of visitors = 136910, number of links = 636497), the Zhu and Ghodsi [49]’s profile-likelihood criterion indicated a six dimensional latent feature space (*d* = 6) as adequate. This accounted for approximately to 70 % variability of the visitation network data (Figure 2).

**Figure 2:**
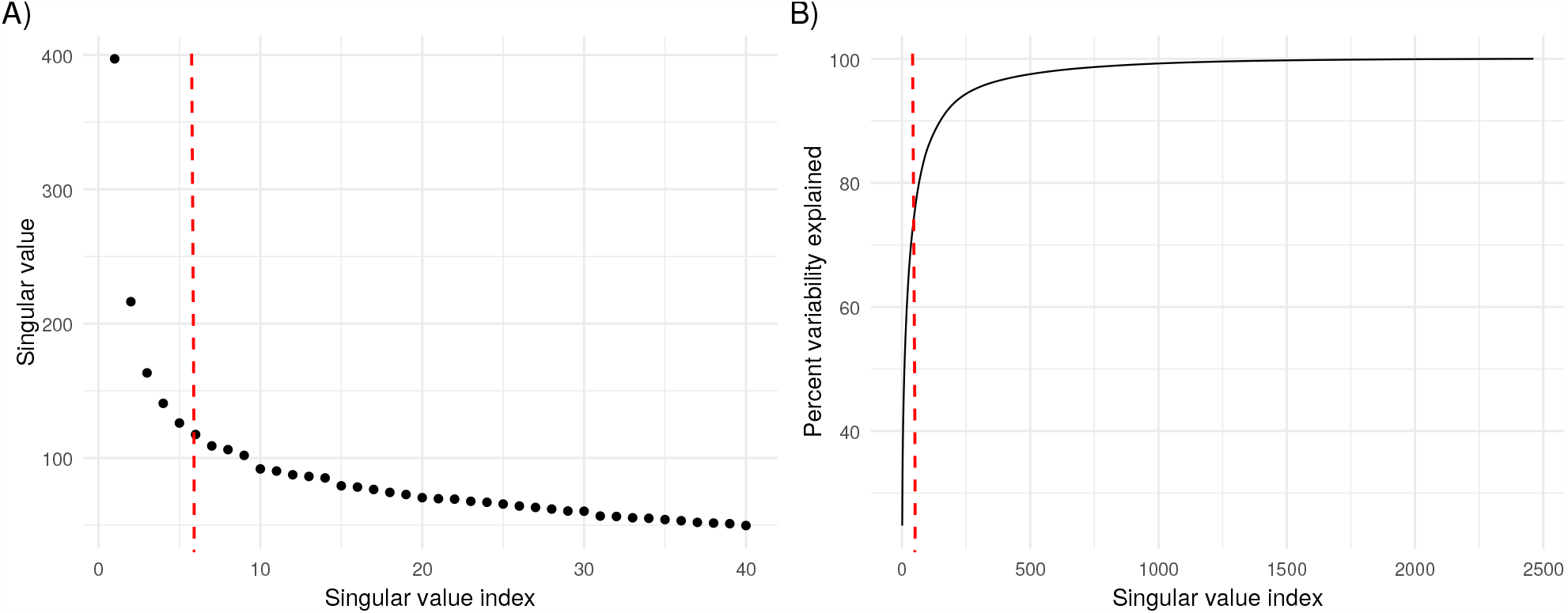
Identifying an adequate dimension *d* of network data. A) The scree plot represents the singular values of the adjacency matrix of the visitation network in decreasing order of *d*. The x-axis shows the singular value index, and the y-axis indicates the singular values. B) Cumulative plot showing the percentage variability explained with the increasing singular value indexes. The x-axis again shows the singular value index and the y-axis indicates the percent of variance explained. Using Zhu and Ghodsi [49]’s likelihood criteria, we picked *d* = 6 as indicated by the red dotted line. This dimension explains 70 % of the variability of the visitation network data.

Overall we found that the neural networks performed better than the linear regression model in finding the best mapping from the node metadata to the latent feature spaces. More specifically, for the visitor metadata, we found that the neural network with two hidden layers (Mean Squared Error (MSE) = 0.0009) and multilayer perceptron performed (MSE = 0.0010) better compared to the linear regression model (MSE = 0.0014) using a constant learning rate (Figure 3). We found similar patterns for the Time-based learning rate (Figure S1).

**Figure 3:**
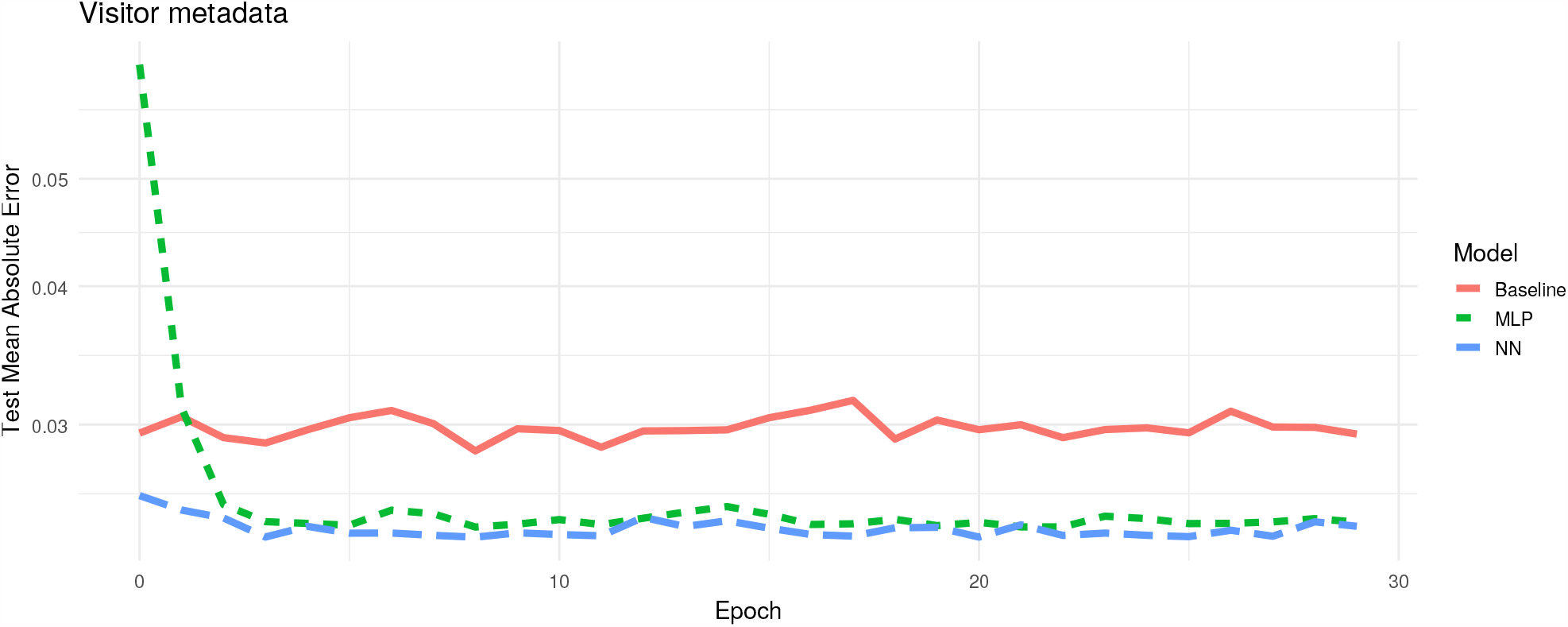
Training of regression models over time when projecting observed visitor metadata onto the latent feature space using an adaptive moment estimation (Adam) optimiser run with 30 epochs and a batch size of 20. The plot shows the model validation—i.e. the subset of training visitation dataset used—to validate the three different models in finding the best mapping from the node metadata to the latent feature space. The x-axis indicates the epochs. The y-axis indicates the Mean Absolute Error (MAE), which is the cost function used to measure the accuracy of model predictions—i.e. it measures the distance between the estimated latent feature space (SVD) and the predicted latent feature space. The red line shows the learning rate of the linear regression model (Baseline), the green line indicates the learning rate of the multilayer perceptron model (MLP), and the blue line indicates the neural network with two hidden layers (NN). The plot shows that both the MLP model and NN model performed better than the Baseline model.

For the place metadata, we found that the the multilayer perceptron model (MSE = 0.125) and linear regression model (MSE = 0.141) performed better than the neural network with two hidden layers (MSE = 0.143) (Figure 4). We also observed similar patterns for models run with the Time-based learning rate (Supplementary Information).

**Figure 4:**
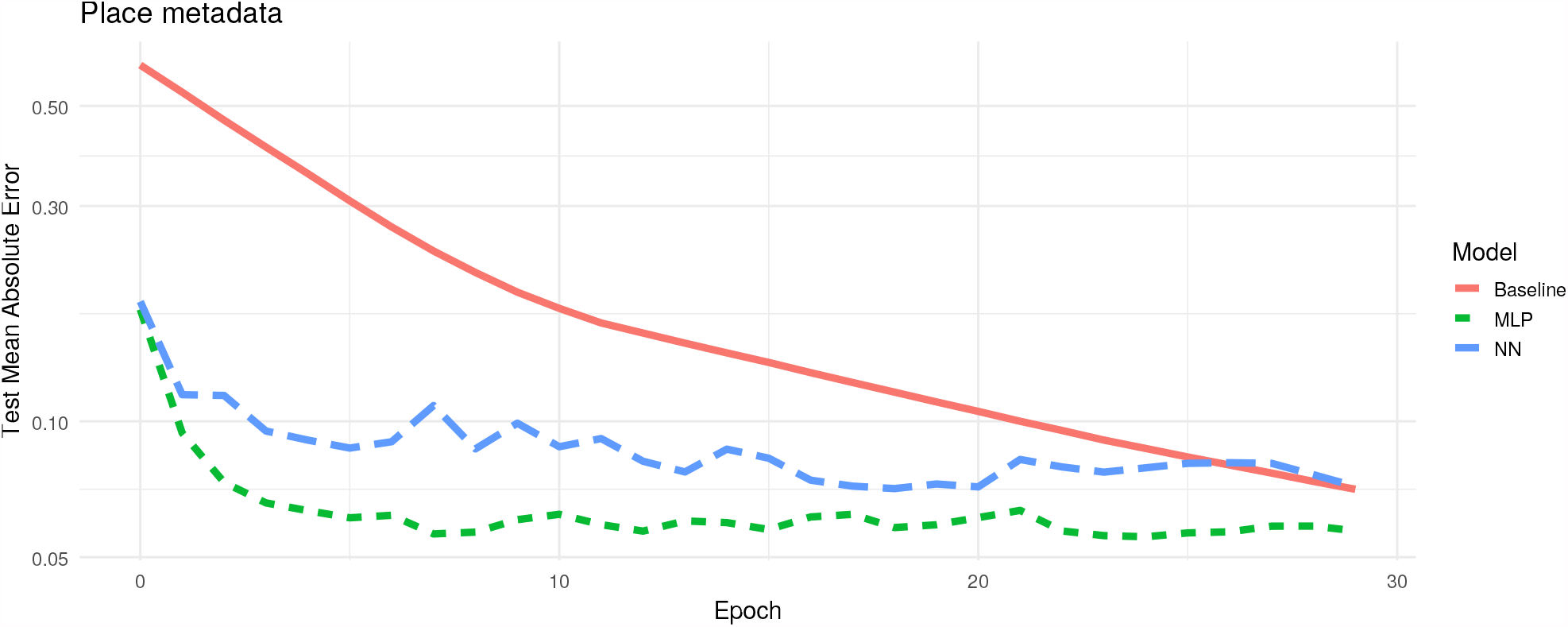
Training of regression models over time when projecting observed place metadata onto the latent feature space using an adaptive moment estimation (Adam) optimiser run with 30 epochs and a batch size of 20. The plot shows the model validation—i.e. the subset of training visitation dataset used—to validate the three different models in finding the best mapping from the node metadata to the latent feature space. The x-axis indicates the epochs. The y-axis indicates the Mean Absolute Error (MAE), which is the cost function used to measure the accuracy of model predictions—i.e. it measures the distance between the estimated latent feature space (SVD) and the predicted latent feature space. The red line shows the learning rate of linear regression model (Baseline), the green line indicates the learning rate of the multilayer perceptron model (MLP), and the blue line indicates the neural network with two hidden layers (NN). Here the MLP model seems to perform better than the baseline and NN models.

The predictive procedure we proposed performed significantly better than at random (Table S5). Comparing the different latent feature prediction models, we found that the dot product of the visitor linear regression model and the place MLP (model 2: neural network with one hidden layer) performed better with *AUC* = 0.736, followed by dot product of the visitor MLP and the place MLP model with *AUC* = 0.701 (Table 3).

**Table 3:**
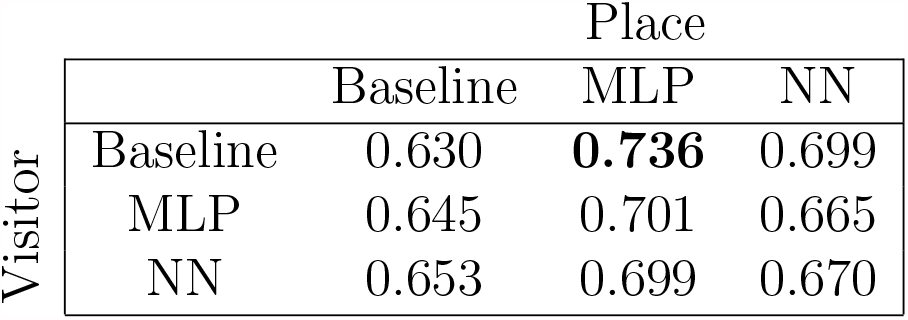
Accuracy of model predictions obtained from RDPG-regression procedure. The table indicates the Area Under Curve (AUC) values for each model calculated using Mean Absolute Error as the cost function to measure the distance between the estimated latent feature spaces and the predicted latent feature space.

| Visitor | Place |  |  |  |
| --- | --- | --- | --- | --- |
|  |  | Baseline | MLP | NN |
|  | Baseline | 0.630 | **0.736** | 0.699 |
|  | MLP | 0.645 | 0.701 | 0.665 |
|  | NN | 0.653 | 0.699 | 0.670 |

## Discussion

In the current study, we present a new predictive procedure which allows us to use both the nodes’ metadata and the knowledge gained from the observed network to predict interactions in large scale networks. More specifically, we showed how to extend the RDPG model—which is a graph embedding method—with a combination of statistical and machine learning techniques. Doing so allowed us to directly relate the nodes’ metadata to their corresponding latent feature spaces. To further illustrate the application of the presented procedure on real world data, we used a large data set of tourist travelling patterns within New Zealand. Overall, we showed that the predictive procedure works in a real-world context with an accuracy of *AUC* = 0.736, which indicates that the procedure performed better than at random. Moreover, we showed that our procedure also allowed us to predict interactions for new nodes added to the network.

To our knowledge, few studies have focused on exploiting the nodes’ metadata to predict interactions using graph-embedding methods. Most research including node metadata to predict interactions have used node-aggregating methods [26, 40, 41]. However, such approaches assume that all nodes belonging to a given group behave identically, ignoring that certain nodes within the given group might be interacting with other nodes in the network to different extents. We instead propose using a predictive procedure based on the RDPG model combined with a machine learning algorithm to account for the heterogeneity of interactions observed at the node level. Moreover, using the truncated SVD allows us to represent an observed network at a lower dimension, which simplified the task of directly relating the observed nodes’ metadata to the estimated latent feature spaces (obtained from the truncated SVD). Using the inferred relationship of nodes’ metadata and the estimated latent feature spaces from the neural networks, we can predict the position of both observed and new nodes in a predicted latent feature space using only the nodes’ metadata and knowledge form the observed network. Then, using the statistical properties of the RDPG model, we can proceed to predict interactions at the node level in a network by simply calculating the dot product of the given node to the other nodes present in the network [4,25,37].

Machine learning techniques such as neural networks are increasingly popular tools in various applications due to their high predictive accuracy [29,46]. Here, we used neural networks to relate the nodes’ metadata to their latent feature spaces obtained from a truncated SVD. Various studies suggest that deeper neural networks—i.e. neural networks with a high number of hidden layers—tend to outperform shallow neural network in a wide variety of tasks [12,16]. While both of the neural network architectures we tested outperformed the linear regression model in mapping the nodes’ metadata onto the latent feature spaces, our results showed that the linear regression model (for the visitors’ metadata) and the neural network with one hidden layer (for the places’ metadata) outperformed the the neural network with two hidden layers in predicting links. This therefore suggests that simpler models can outperform deeper neural networks in at least some situations. Note, however, that the main purpose of our study was not to find the absolute best neural network architecture, and hence we do not expect further studies to necessarily confirm this result.

Metadata are known to be good proxies from which to predict interactions in a network [17, 30, 32, 33, 41]. However, node metadata can be of varied type—i.e. categorical and continuous variables, and these variables might not have linear relationships to the latent feature space; these factors together necessitate different modelling frameworks [3,26]. Here, we showed that the high flexibility of neural networks (or other machine learning algorithms) enabled the identification of an accurate mapping from the visitors’ metadata onto the visitors’ latent feature space and from the places’ metadata to the places’ latent feature space, respectively. As the functional relationship between nodes’ metadata and their position in the latent feature space can be very complicated, neural network methods are a promising approach to learn it.

While each step of the presented procedure is robust, there might be many sources of error. In the current study, we only present an exploratory analysis of a bipartite network to predict interactions in a visitation network using both node types’ metadata. Rather than attempting to find the optimal dimension of our network data, we instead chose *d* according to the Zhu and Ghodsi [49]’s profile-likelihood criterion. We selected *d a priori*, based only on the topological structure of the network. It could be interesting to further explore whether using a different procedure to select the dimension of the latent space improves the accuracy of link prediction. Indeed, *a posteriori* selection (trying to identify which dimension *d* grants a higher prediction accuracy) is another possibility, but may require substantially greater computational effort.

The way in which we split our training and test set implies that the observed and new nodes’ metadata are sampled from the same distribution. Therefore, the models learn a relevant mapping of the new nodes into a suitable region of the latent feature space. However, if this is not the case, and the metadata of the new nodes is completely different from the one of the observed nodes, nothing guarantees a good placement in the latent feature space. How to deal with new nodes with “surprising” metadata is an open problem.

In addition, we assumed all the nodes’ metadata could be informative when predicting interactions. As a result, we learnt a mapping from the nodes’ full metadata to their respective latent feature spaces obtained from the truncated SVD. However, we know that not all of the metadata is necessarily informative, specially when predicting interactions [11,32]. Therefore, further investigating the relationship between the nodes’ metadata and the latent feature space obtained from the truncated SVD should be done to understand whether certain node metadata are affecting the link prediction accuracy in the presented procedure.

Moving forward, it would be interesting to extend the current procedure to account for missing data in: 1) the interaction probability matrix—i.e. distinguishing new and absent interactions—and 2) the metadata—i.e. when some of the nodes’ metadata are missing. One can imagine a scenario where a survey was carried out, and a person did not complete the full survey. If we were to have the metadata of that particular person, we could potentially interpolate some of their answers. Similarly, in the case where data is extracted from an experimental set up, data might be missing as a result of failed experiments. Accounting for such missing information can be particularly important. In this direction, deeper or dedicated neural network architectures, such as the ones in Smieja et al. [39] and Przewikeźlikowski et al. [34], could be used. More recently, Lerique et al. [21] have used a neural network approach to find the joint embedding of metadata and the network structure to predict the interaction probabilities. However, one the main limitations of the latter approach is the need to find an optimal dimension for the both the nodes’ metadata and the network data. Using a machine learning approach to learn a mapping of nodes’ metadata directly to their interaction probabilities in large networks remains a hard problem when performed in a very high dimensional space. Here we showed that we can simplify that problem by exploiting the properties of a well understood statistical model for complex networks, the Random Dot Product Graph model, and combining it with standard machine learning techniques. The RDPG model grants us a robust estimation of a low dimensional network embedding (the nodes’ latent feature spaces) and a convenient way to estimate its dimension. As in other examples [36], promising results are obtained not by abandoning a model-based approach to science but by merging it with machine learning techniques.

## Data Accessibility

All visitation data from the Ministry of Business Innovation & Employment (MBIE) used in this study are publicly available on https://www.mbie.govt.nz/.

## Authors’ contributions

RR and GVDR designed the study. RR contributed to the code, carried out the analysis and wrote the first draft of the manuscript. All authors contributed to framing the manuscript, editing and approving the final draft.

## Acknowledgements

RR and DBS acknowledge funding from the New Zealand’s Biological Heritage Ng—a Koiora Tuku Iho National Science Challenge. The authors would also like to thank Michelle Marraffini, Hao Ran Lai, Stephen Merry, and current members of the Stouffer Lab and DaRe group for feedback and valuable discussions.

## Supplementary Information

### Visitation data

**Table S1:** Summary of visitation data extracted from national surveys

| Organisation | Survey | Years | Places | Visitors | Visitor-Place interactions |
| --- | --- | --- | --- | --- | --- |
| MBIE | IVS | 1997 - 2016 | 548 | 110 750 | 89 309 500 |
| MBIE | DTS | 1997 - 2008 | 2 426 | 84 891 | 473 070 000 |

To get an overview of the visitors’ travelling patterns across New Zealand, we extracted data from three national surveys conducted by the New Zealand Ministry of Business, Innovation and Employment (MBIE) and Department of Conservation (DOC): The International Visitor Survey [28], the Domestic Travel Survey [27] and the National Survey of New Zealanders [10] (Table 1). The International Visitor Survey (IVS) targets international visitors departing New Zealand at the 4 main international airports (Auckland, Wellington, Christchurch and Queenstown) whereas the Domestic Travel Survey (DTS) contacts the domestic travellers via phone interviews about their recent trips. Both surveys record detailed information about the trips travelled by each visitor.

Note that as in the current study we focused on testing our predictive framework, we selected *only* visitors and places with the complete set of information. We therefore resulted with 189942 visitors (out of 195641) and 2616 places (out of 2974).

**Table S2:** Summary of node metadata used for neural networks

| Node type | Metadata | Data type | Output type |
| --- | --- | --- | --- |
| Visitor | Age | Categorical | Single label with multiple classes |
| Visitor | Gender | Categorical | Binary |
| Visitor | Activity type | Categorical | Multiple labels with multiple classes |
| Visitor | Mode of transportation | Categorical | Single label with multiple classes |
| Place | Place geolocation | Continuous | Numerical values |
| Place | Place type | Categorical | Multiple labels with multiple classes |
| Place | Regional Council | Categorical | Single label with multiple classes |

### Mapping node metadata to latent feature spaces

**Table S3:** Regression models used to predict position of nodes using observed metadata

| Metadata | Model |
| --- | --- |
| Visitor | $x_n = \beta_0 + \beta_{Age}x_1 + \beta_{Gender}x_2 + \beta_{Activities}x_3 + \beta_{Transport}x_4$ |
| Place | $y_m = \beta_0 + \beta_{Region}y_1 + \beta_{Placetype}y_2 + \beta_{Latitude}y_3 + \beta_{Longitude}y_4$ |

**Figure S1:**
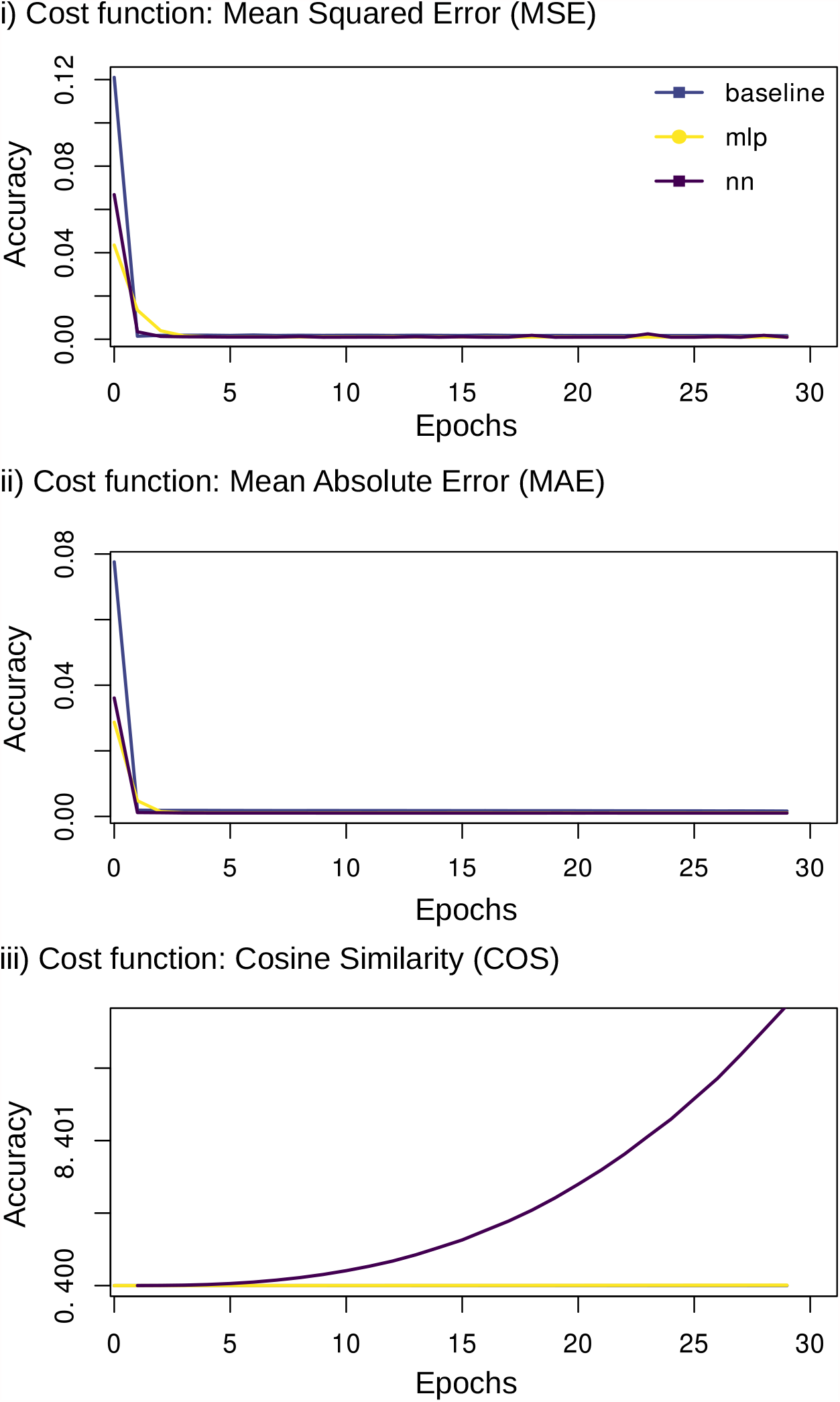
Accuracy of model training when projecting visitor metadata onto the latent feature space using adaptive moment estimation (Adam) optimiser with 100 epochs and a batch size of 20. Here, exploratory analysis are carried out using the Time-based decay learning rate—where the initial learning rate (0.01) decrease by 0.0001 after each epoch. Note that we also used three different cost functions during the model training: Mean Squared Error (MSE), Mean Absolute Error (MAE) and Cosine Similarity (COS) to measure the distance between the estimated latent feature space (truncated SVD) and the predicted latent feature space obtained from the regression models.

**Figure S2:**
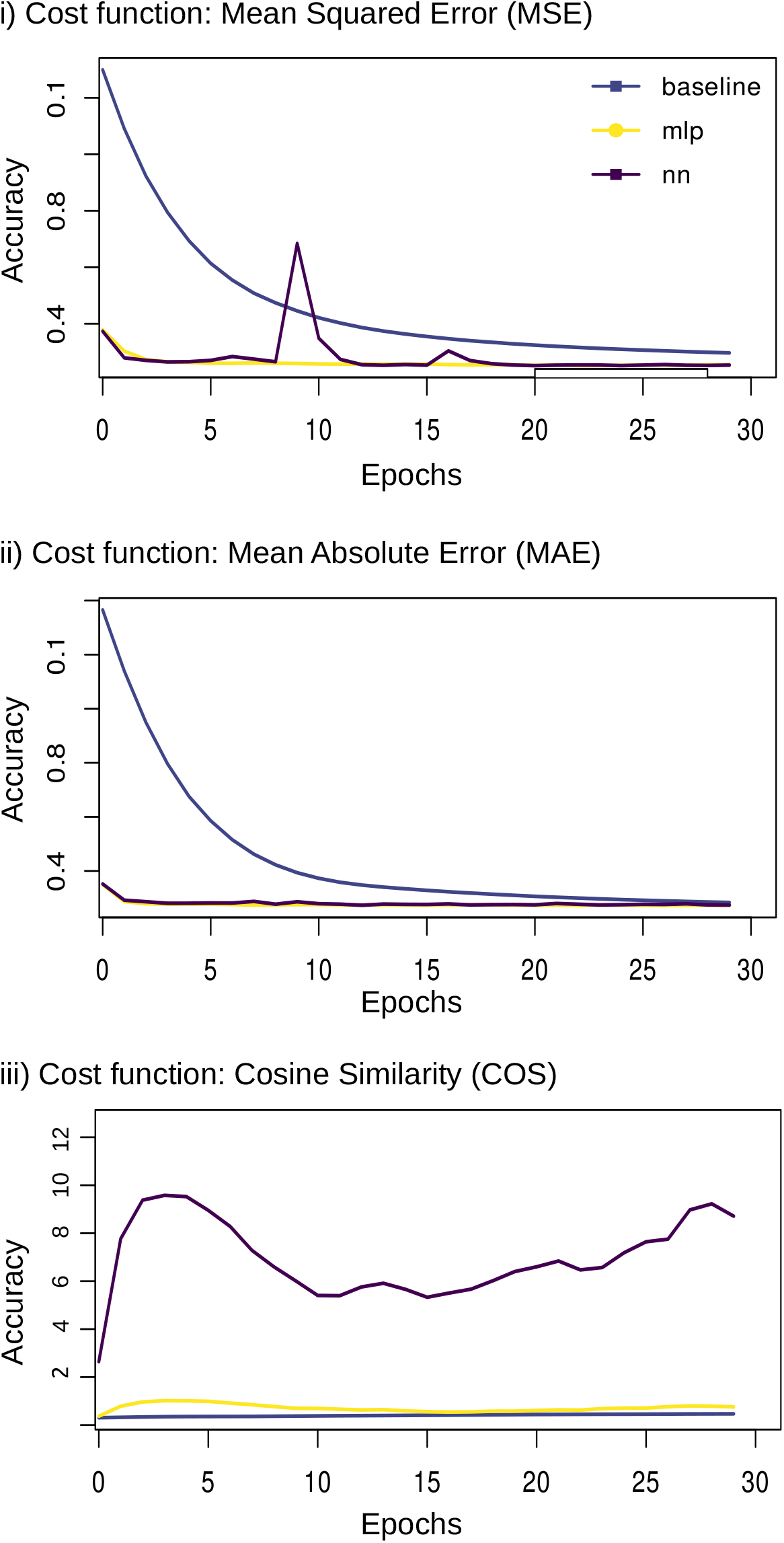
Accuracy of model training when projecting place metadata onto the latent feature space using adaptive moment estimation (Adam) optimiser with 100 epochs and a batch size of 20. Here, exploratory analysis are carried out using the Time-based decay learning rate—where the initial learning rate (0.01) decrease by 0.0001 after each epoch. Note that we also used three different loss functions during the model training: Mean Squared Error (MSE), Mean Absolute Error (MAE) and Cosine Similarity (COS) to measure the distance between the estimated latent feature space (truncated SVD) and the predicted latent feature space obtained from the regression models.

### Predicting interaction using observed node metadata

**Table S4:** List of model predictions obtained using the dot product of inferred latent feature spaces (obtained from regression models)

| Model no. | Visitor | Place | Learning rate |
| --- | --- | --- | --- |
| 1A | Baseline | Baseline | Constant |
| 1B | Baseline | Baseline | Time-based |
| 2A | Baseline | MLP | Constant |
| 2B | Baseline | MLP | Time-based |
| 3A | Baseline | NN | Constant |
| 3B | Baseline | NN | Time-based |
| 4A | MLP | Baseline | Constant |
| 4B | MLP | Baseline | Time-based |
| 5A | MLP | MLP | Constant |
| 5B | MLP | MLP | Time-based |
| 6A | MLP | NN | Constant |
| 6B | MLP | NN | Time-based |
| 7A | NN | Baseline | Constant |
| 7B | NN | Baseline | Time-based |
| 8A | NN | MLP | Constant |
| 8B | NN | MLP | Time-based |
| 9A | NN | NN | Constant |
| 9B | NN | NN | Time-based |

**Table S5:** Accuracy of model predictions obtained from RDPG-regression framework. The table indicates the Area Under Curve (AUC) values for each model calculated using three different metrics which was used to measure the distance between the estimated latent feature spaces and the predicted latent feature space: Mean Squared Error (MSE), Mean Absolute Error (MAE) and Cosine Similarity (COS).

| Model | MSE | MAE | COS |
| --- | --- | --- | --- |
| 1A | 0.495 | 0.630 | 0.511 |
| 1B | 0.530 | 0.538 | **0.729** |
| 2A | **0.733** | **0.736** | 0.509 |
| 2B | 0.584 | 0.699 | 0.574 |
| 3A | 0.584 | 0.699 | 0.574 |
| 3B | 0.610 | 0.636 | 0.516 |
| 4A | 0.712 | 0.645 | 0.510 |
| 4B | 0.620 | 0.576 | 0.720 |
| 5A | 0.668 | 0.701 | 0.532 |
| 5B | 0.550 | 0.646 | 0.516 |
| 6A | 0.620 | 0.665 | 0.610 |
| 6B | 0.576 | 0.590 | 0.516 |
| 7A | 0.632 | 0.653 | 0.515 |
| 7B | 0.502 | 0.615 | 0.719 |
| 8A | 0.721 | 0.699 | 0.488 |
| 8B | 0.537 | 0.657 | 0.499 |
| 9A | 0.500 | 0.670 | 0.570 |
| 9B | 0.601 | 0.604 | 0.530 |

**Figure S3:**
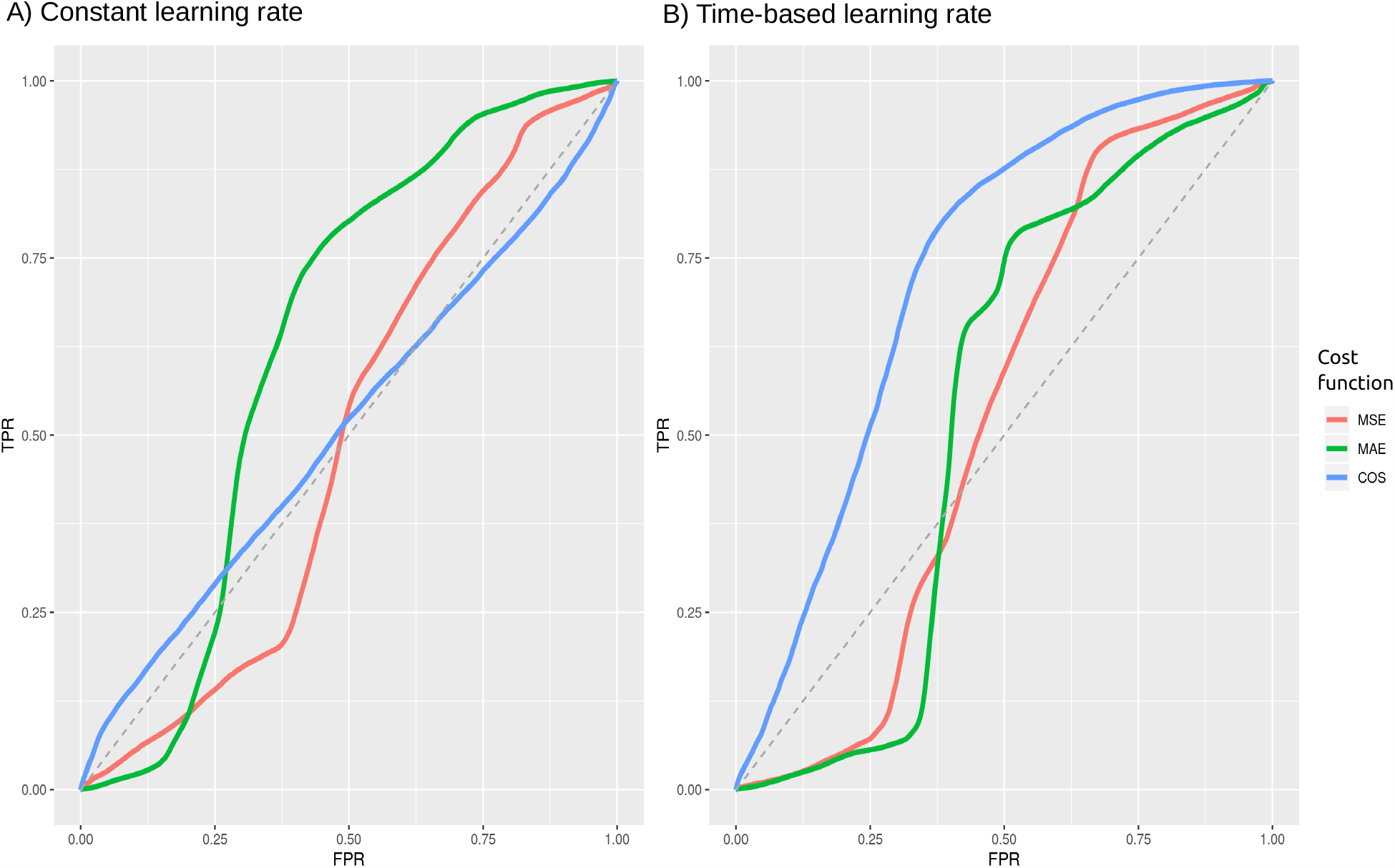
Measuring the performance of predictive framework using the AUC-ROC. The plot shows the accuracy of predictions obtained from the dot product in predicting interactions for the model 1 (dot product of visitor baseline and place baseline models) using A) Constant learning rate and B) Time-based learning rate. Note that rate of recovering False Positive and True Positive are both calculated at thresholds varying from 0 to 1. The dotted line indicates an *AUC* = 0.5 where a model does not distinguish between True Positives and False Positives. We also report the model predictions obtained using different metrics: Mean Squared Error (MSE -in red), Mean Absolute Error (MAE -in green) and Cosine Similarity (COS -in blue) which measures the distance between the estimated latent feature space and the predicted latent feature spaces.

**Figure S4:**
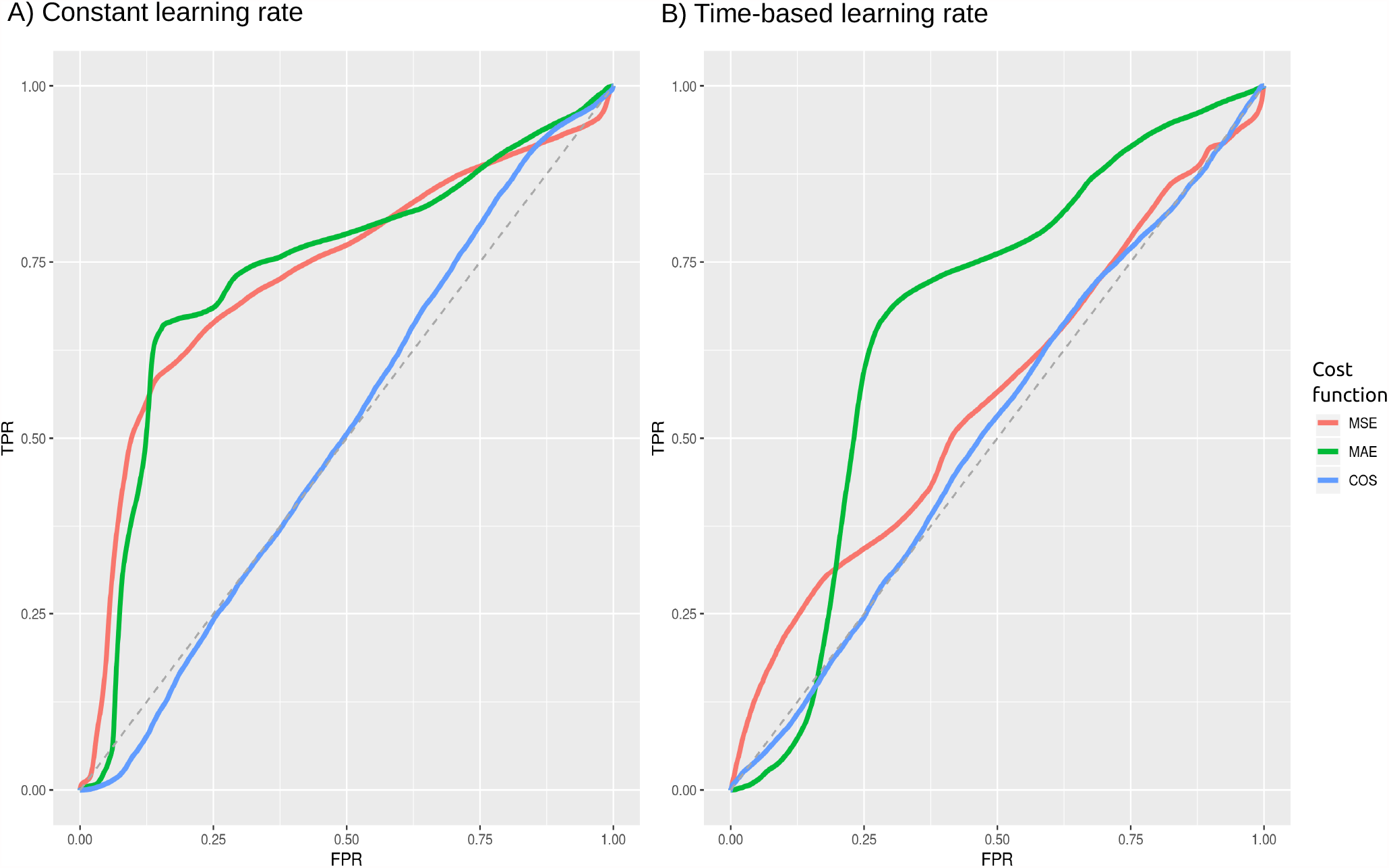
Measuring the performance of predictive framework using the AUC-ROC. The plot shows the accuracy of predictions obtained from the dot product in predicting interactions for the model 2 (dot product of visitor Baseline and place MLP models) using A) Constant learning rate and B) Time-based learning rate. The x-axis indicates the False Positive Rate (FPR) and y-axis indicates the True Positive Rate (TPR). Note that rate of recovering False Positive and True Positive are both calculated at thresholds varying from 0 to 1. The dotted line indicates an *AUC* = 0.5 where a model does not distinguish between True Positives and False Positives. We also report the model predictions obtained using different metrics: Mean Squared Error (MSE -in red), Mean Absolute Error (MAE -in green) and Cosine Similarity (COS -in blue) which measures the distance between the estimated latent feature space and the predicted latent feature spaces.

**Figure S5:**
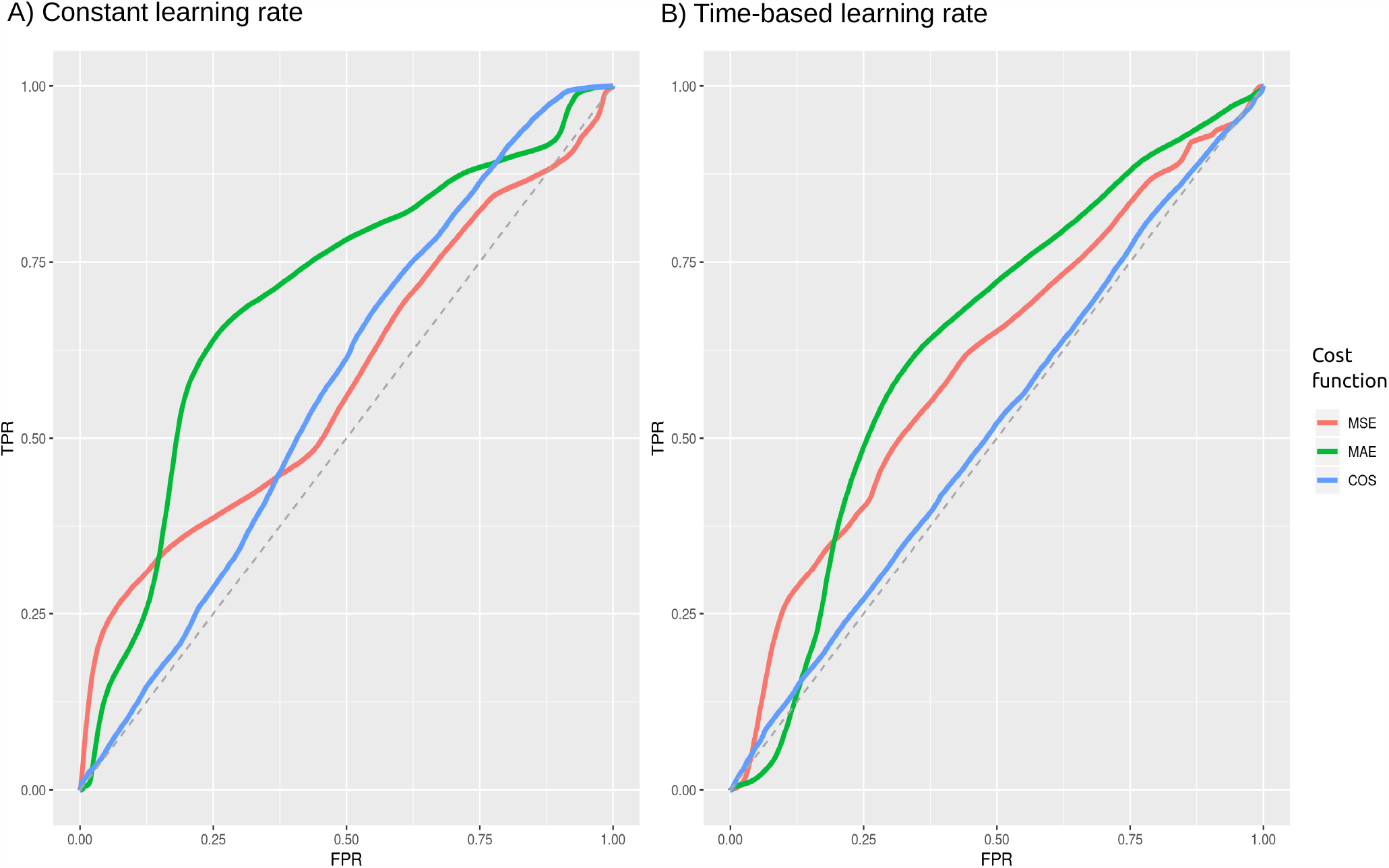
Measuring the performance of predictive framework using the AUC-ROC. The plot shows the accuracy of predictions obtained from the dot product in predicting interactions for the model 3 (dot product of visitor Baseline and place NN models) using A) Constant learning rate and B) Time-based learning rate. The x-axis indicates the False Positive Rate (FPR) and y-axis indicates the True Positive Rate (TPR). Note that rate of recovering False Positive and True Positive are both calculated at thresholds varying from 0 to 1. The dotted line indicates an *AUC* = 0.5 where a model does not distinguish between True Positives and False Positives. We also report the model predictions obtained using different metrics: Mean Squared Error (MSE in red), Mean Absolute Error (MAE -in green) and Cosine Similarity (COS -in blue) which measures the distance between the estimated latent feature space and the predicted latent feature spaces.

**Figure S6:**
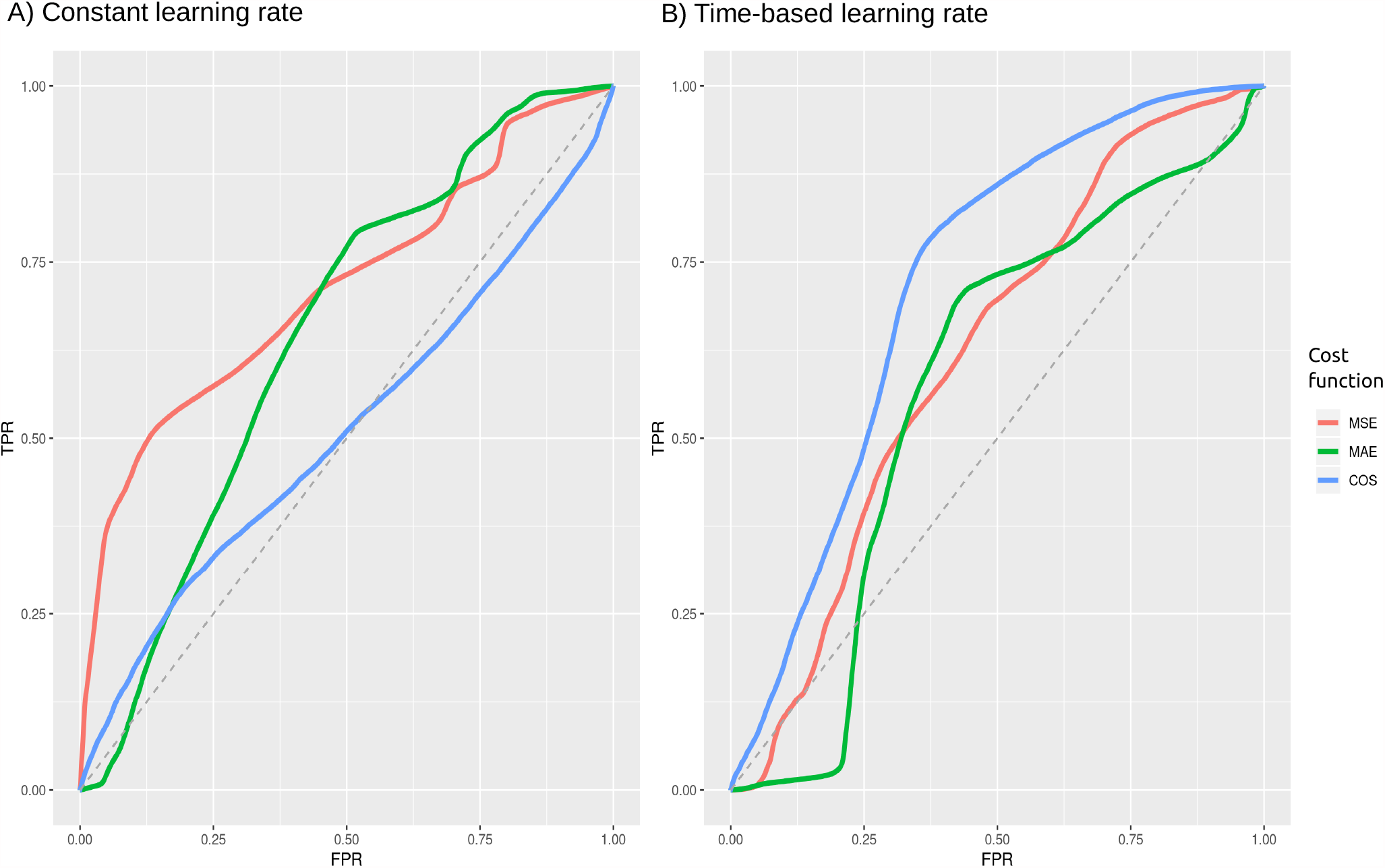
Measuring the performance of predictive framework using the AUC-ROC. The plot shows the accuracy of predictions obtained from the dot product in predicting interactions for the model 4 (dot product of visitor MLP and place Baseline models) using A) Constant learning rate and B) Time-based learning rate. The x-axis indicates the False Positive Rate (FPR) and y-axis indicates the True Positive Rate (TPR). Note that rate of recovering False Positive and True Positive are both calculated at thresholds varying from 0 to 1. The dotted line indicates an *AUC* = 0.5 where a model does not distinguish between True Positives and False Positives. We also report the model predictions obtained using different metrics: Mean Squared Error (MSE -in red), Mean Absolute Error (MAE -in green) and Cosine Similarity (COS -in blue) which measures the distance between the estimated latent feature space and the predicted latent feature spaces.

**Figure S7:**
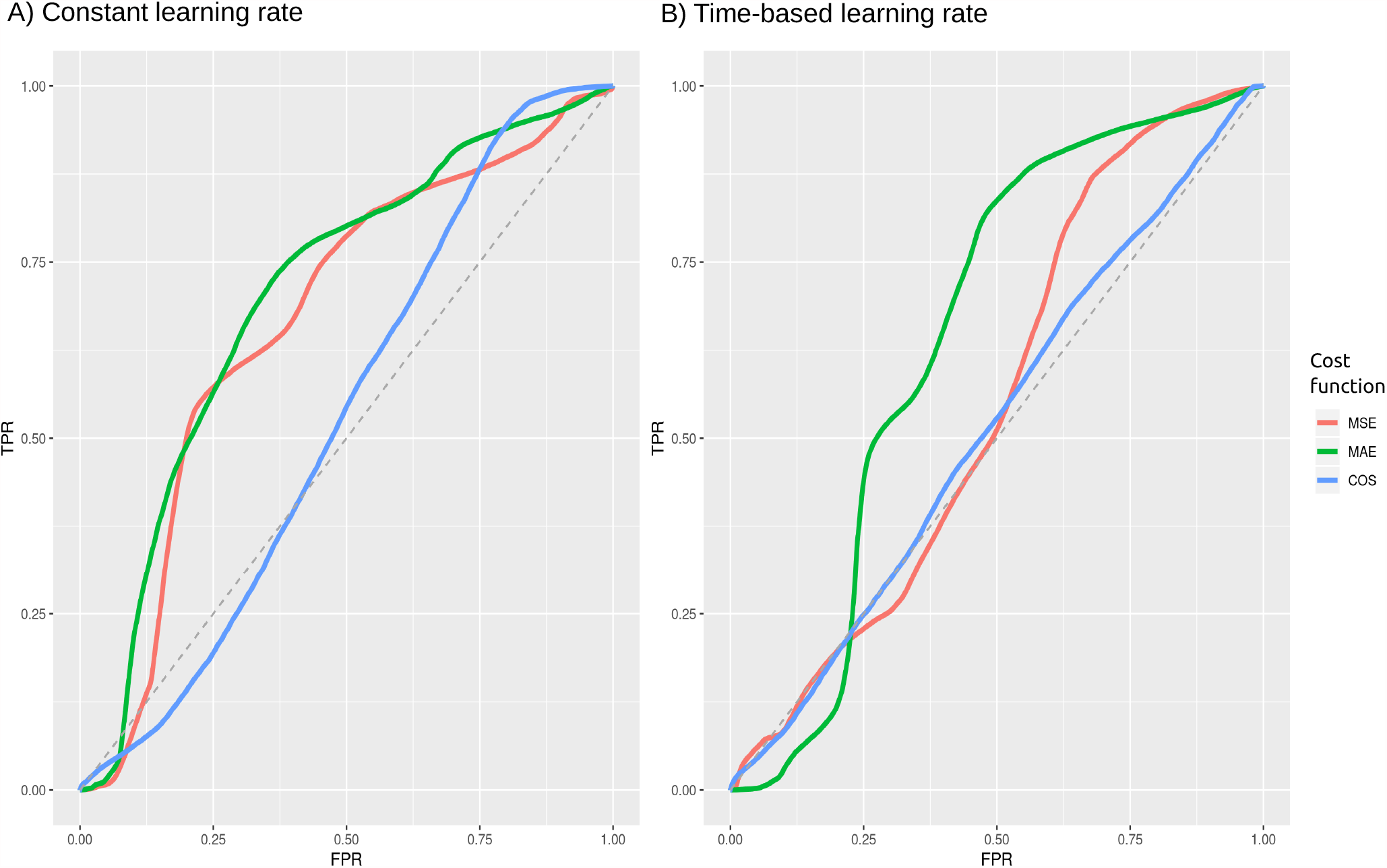
Measuring the performance of predictive framework using the AUC-ROC. The plot shows the accuracy of predictions obtained from the dot product in predicting interactions for the model 5 (dot product of visitor MLP and place MLP models) using A) Constant learning rate and B) Time-based learning rate. The x-axis indicates the False Positive Rate (FPR) and y-axis indicates the True Positive Rate (TPR). Note that rate of recovering False Positive and True Positive are both calculated at thresholds varying from 0 to 1. The dotted line indicates an *AUC* = 0.5 where a model does not distinguish between True Positives and False Positives. We also report the model predictions obtained using different metrics: Mean Squared Error (MSE -in red), Mean Absolute Error (MAE -in green) and Cosine Similarity (COS -in blue) which measures the distance between the estimated latent feature space and the predicted latent feature spaces.

**Figure S8:**
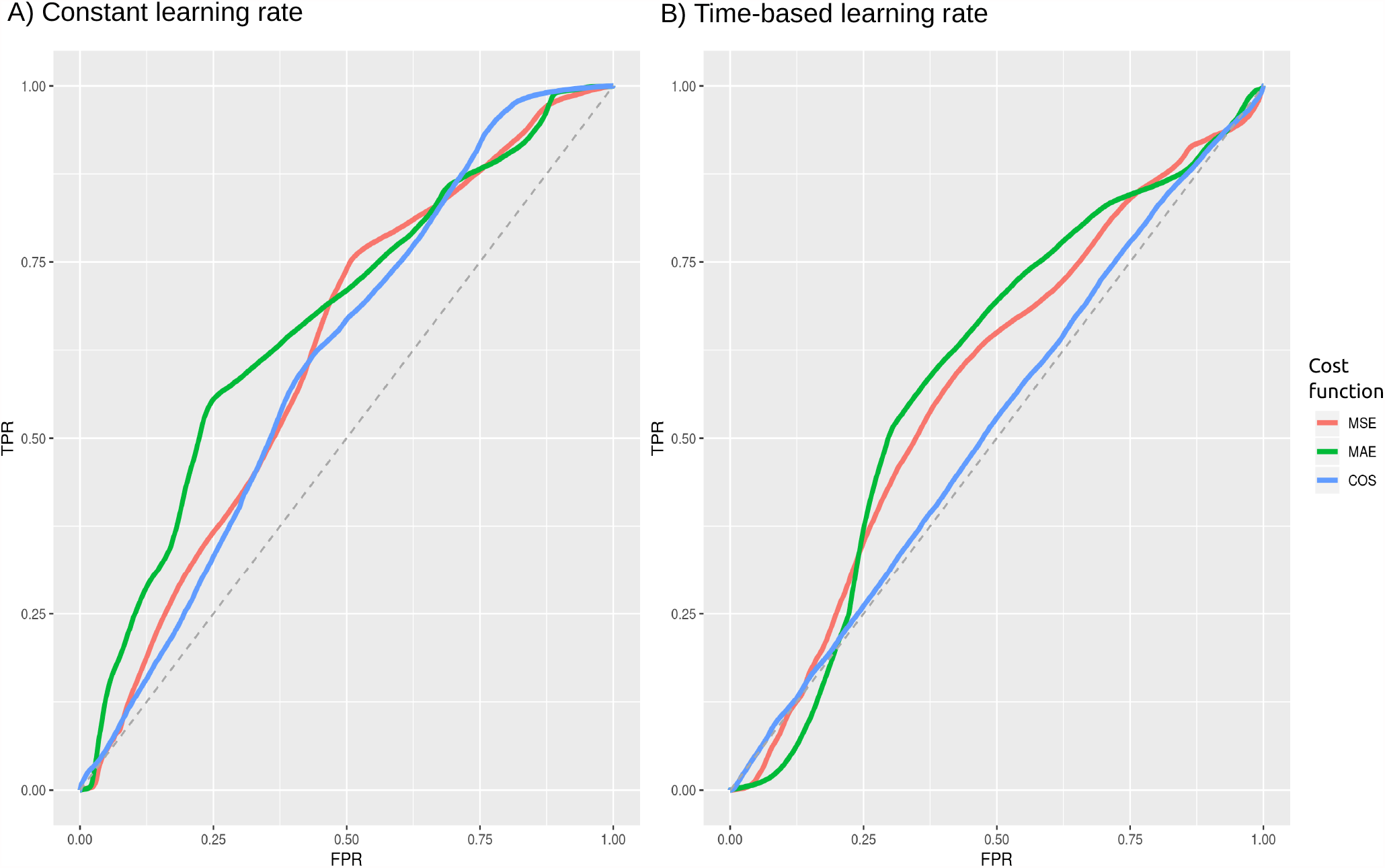
Measuring the performance of predictive framework using the AUC-ROC. The plot shows the accuracy of predictions obtained from the dot product in predicting interactions for the model 6 (dot product of visitor MLP and place NN models) using A) Constant learning rate and B) Time-based learning rate. The x-axis indicates the False Positive Rate (FPR) and y-axis indicates the True Positive Rate (TPR). Note that rate of recovering False Positive and True Positive are both calculated at thresholds varying from 0 to 1. The dotted line indicates an *AUC* = 0.5 where a model does not distinguish between True Positives and False Positives. We also report the model predictions obtained using different metrics: Mean Squared Error (MSE in red), Mean Absolute Error (MAE -in green) and Cosine Similarity (COS -in blue) which measures the distance between the estimated latent feature space and the predicted latent feature spaces.

**Figure S9:**
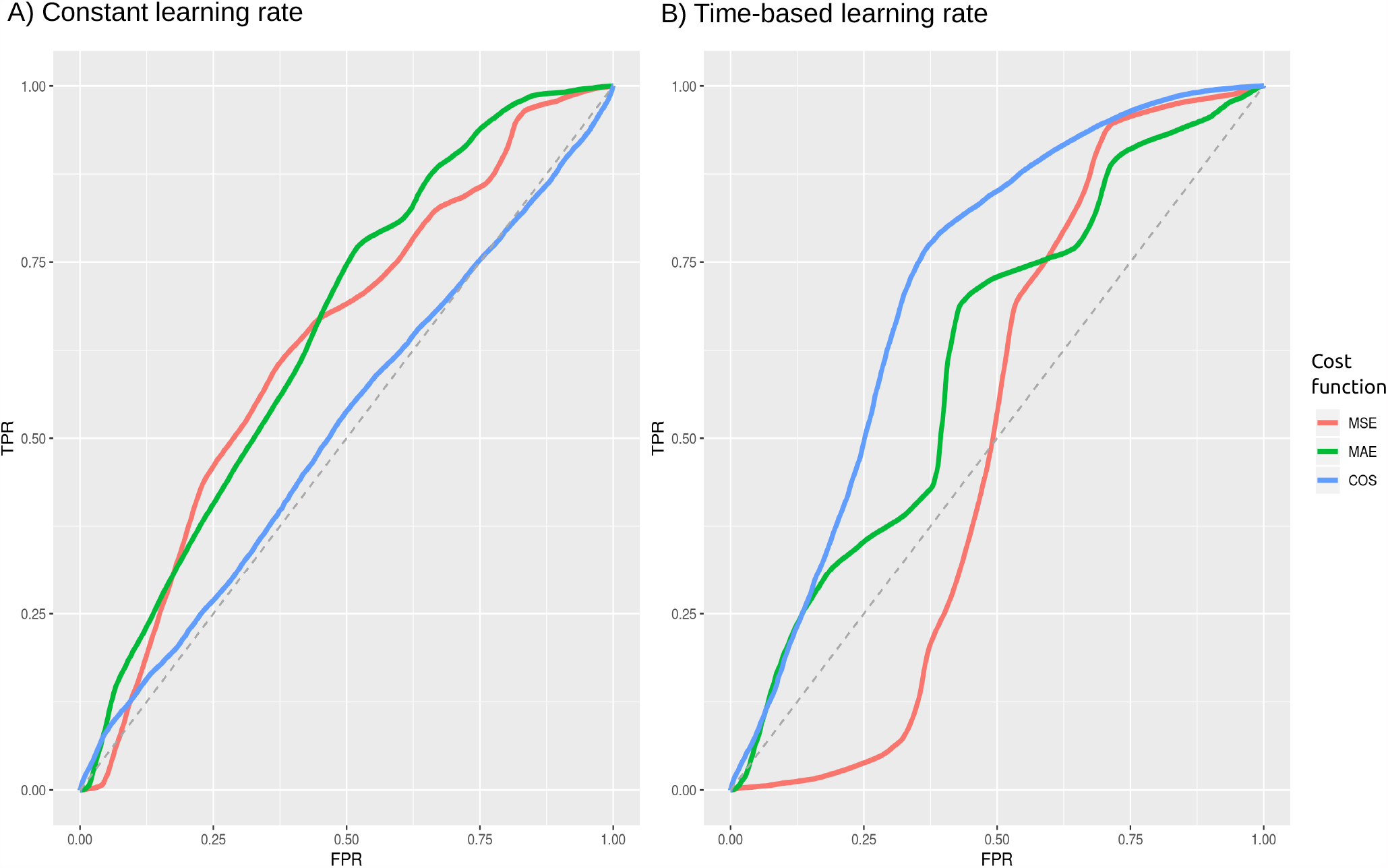
Measuring the performance of predictive framework using the AUC-ROC. The plot shows the accuracy of predictions obtained from the dot product in predicting interactions for the model 7 (dot product of visitor NN and place Baseline models) using A) Constant learning rate and B) Time-based learning rate. The x-axis indicates the False Positive Rate (FPR) and y-axis indicates the True Positive Rate (TPR). Note that rate of recovering False Positive and True Positive are both calculated at thresholds varying from 0 to 1. The dotted line indicates an *AUC* = 0.5 where a model does not distinguish between True Positives and False Positives. We also report the model predictions obtained using different metrics: Mean Squared Error (MSE -in red), Mean Absolute Error (MAE -in green) and Cosine Similarity (COS -in blue) which measures the distance between the estimated latent feature space and the predicted latent feature spaces.

**Figure S10:**
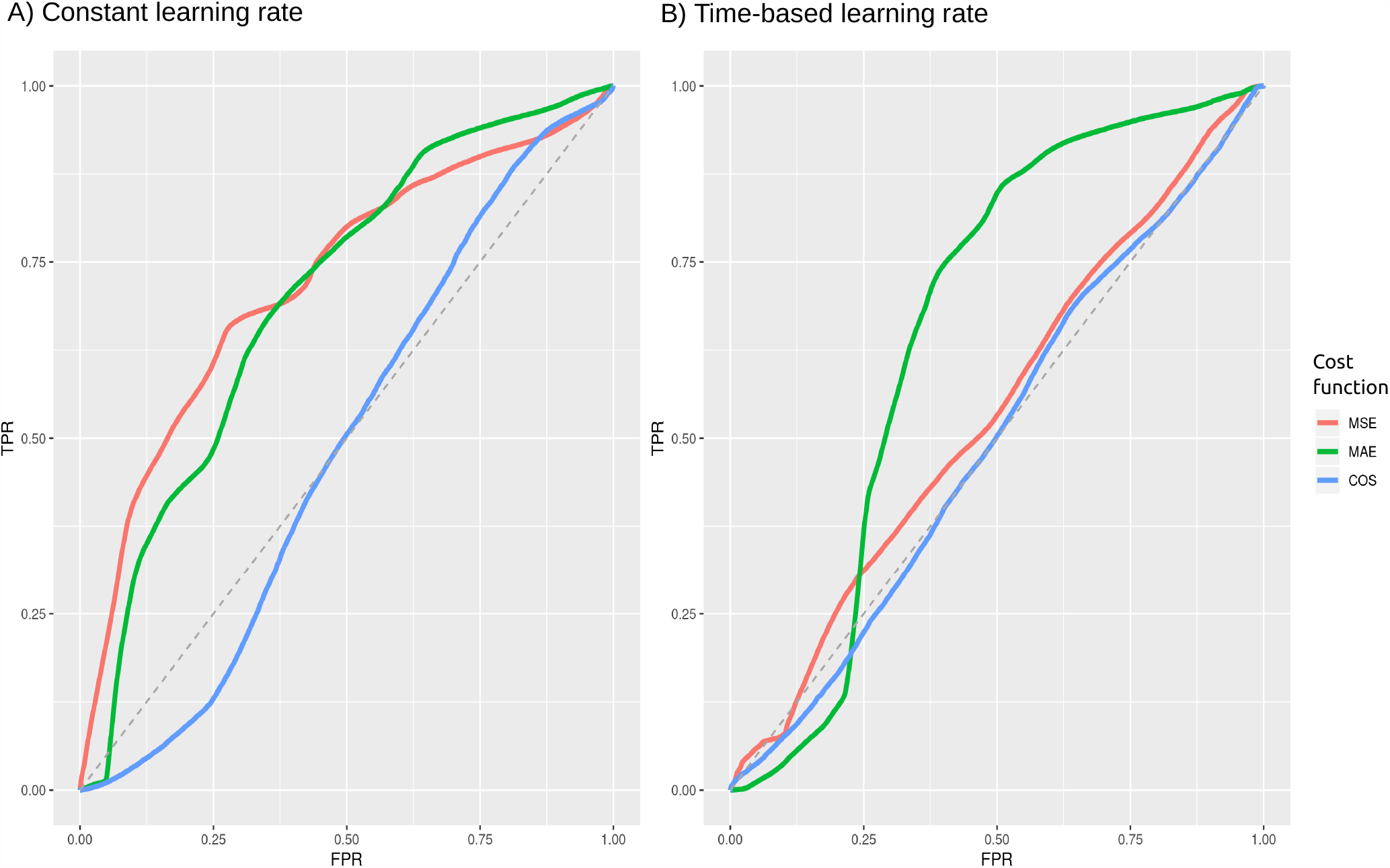
Measuring the performance of predictive framework using the AUC-ROC. The plot shows the accuracy of predictions obtained from the dot product in predicting interactions for the model 8 (dot product of visitor NN and place MLP models) using A) Constant learning rate and B) Time-based learning rate. The x-axis indicates the False Positive Rate (FPR) and y-axis indicates the True Positive Rate (TPR). Note that rate of recovering False Positive and True Positive are both calculated at thresholds varying from 0 to 1. The dotted line indicates an *AUC* = 0.5 where a model does not distinguish between True Positives and False Positives. We also report the model predictions obtained using different metrics: Mean Squared Error (MSE in red), Mean Absolute Error (MAE -in green) and Cosine Similarity (COS -in blue) which measures the distance between the estimated latent feature space and the predicted latent feature spaces.

**Figure S11:**
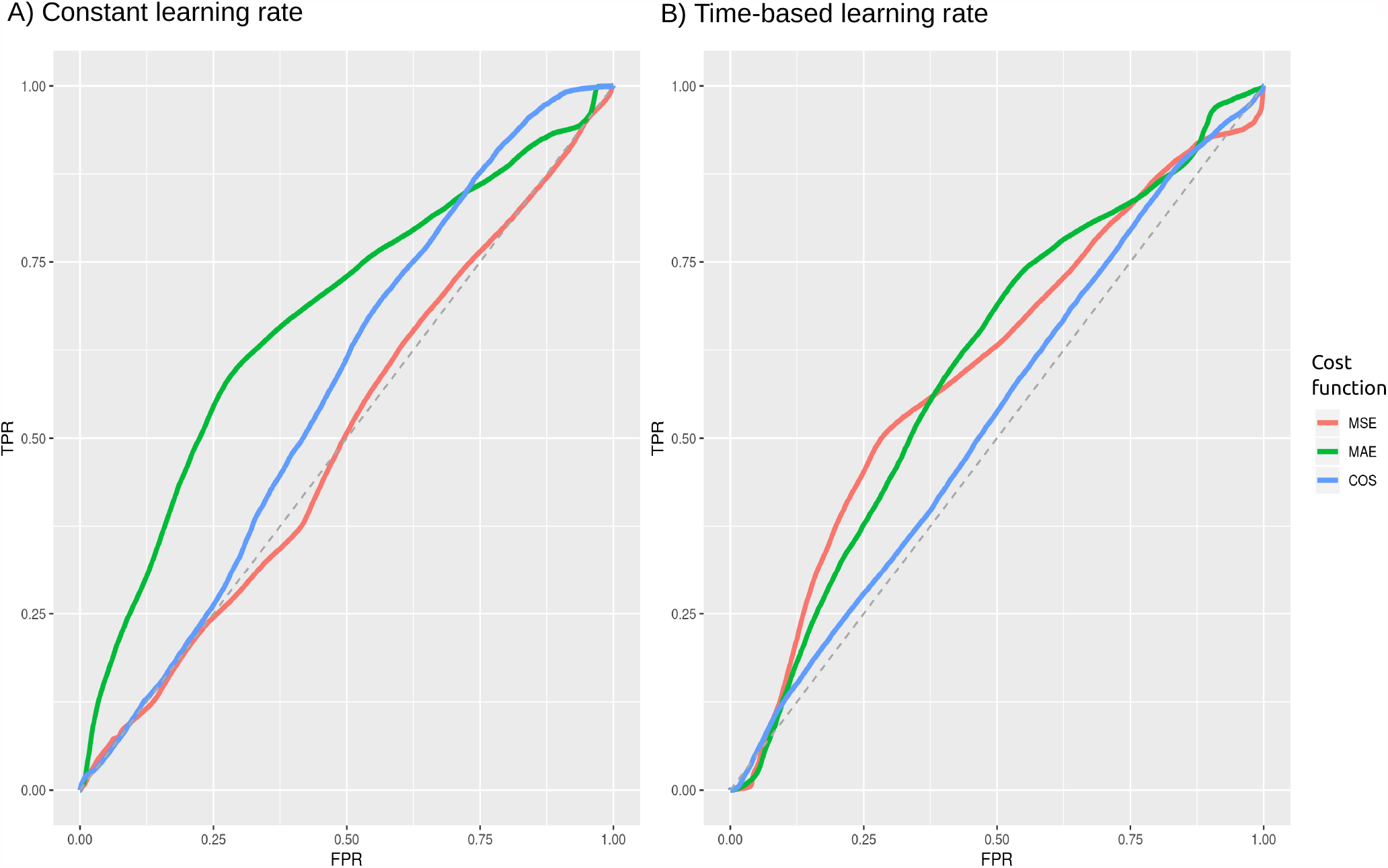
Measuring the performance of predictive framework using the AUC-ROC. The plot shows the accuracy of predictions obtained from the dot product in predicting interactions for the model 9 (dot product of visitor NN and place NN models) using A) Constant learning rate and B) Time-based learning rate. The x-axis indicates the False Positive Rate (FPR) and y-axis indicates the True Positive Rate (TPR). Note that rate of recovering False Positive and True Positive are both calculated at thresholds varying from 0 to 1. The dotted line indicates an *AUC* = 0.5 where a model does not distinguish between True Positives and False Positives. We also report the model predictions obtained using different metrics: Mean Squared Error (MSE in red), Mean Absolute Error (MAE -in green) and Cosine Similarity (COS -in blue) which measures the distance between the estimated latent feature space and the predicted latent feature spaces.

